# SARS-CoV-2 Genome Sequencing Methods Differ In Their Ability To Detect Variants From Low Viral Load Samples

**DOI:** 10.1101/2021.05.01.442304

**Authors:** C. Lam, K. Gray, M. Gall, R. Sadsad, A. Arnott, J. Johnson-Mackinnon, W. Fong, K. Basile, J. Kok, D. E. Dwyer, V. Sintchenko, R.J. Rockett

## Abstract

SARS-CoV-2 genomic surveillance has been vital in understanding the spread of COVID-19, the emergence of viral escape mutants and variants of concern. However, low viral loads in clinical specimens affect variant calling for phylogenetic analyses and detection of low frequency variants, important in uncovering infection transmission chains. We systematically evaluated three widely adopted SARS-CoV-2 whole genome sequencing methods for their sensitivity, specificity, and ability to reliably detect low frequency variants. Our analyses highlight that the ARTIC v3 protocol consistently displays high sensitivity for generating complete genomes at low viral loads compared with the probe-based Illumina respiratory viral oligo panel, and a pooled long-amplicon method. We show substantial variability in the number and location of low-frequency variants detected using the three methods, highlighting the importance of selecting appropriate methods to obtain high quality sequence data from low viral load samples for public health and genomic surveillance purposes.

## INTRODUCTION

The rapid implementation of genomic epidemiology has enabled unparalleled understanding and monitoring of viral evolution during the SARS-CoV-2 pandemic. The first reported SARS-CoV-2 case in Australia was notified on January 25, 2020, and by the end of 2020, 28,381 SARS-CoV-2 cases had been identified nationwide (https://www.health.gov.au/resources/publications/coronavirus-covid-19-at-a-glance-30-december-2020-0). Australia’s low prevalence of COVID-19 is due to the implementation of strong public health measures, which in New South Wales (NSW) has included integrated genomic surveillance to inform public health responses and contact tracing efforts [1].

Whole genome sequencing (WGS) of SARS-CoV-2 was implemented in NSW, Australia within two weeks of the first reported case in anticipation of increasing SARS-CoV-2 infections [2]. A pooled long-amplicon (long-amp) based sequencing approach was initially selected based on reagent and resource availability and was quickly adapted to fit existing WGS workflows and infrastructure [3]. By March 28, 2020, 209 samples from NSW had been sequenced and released on the Global Initiative on Sharing All Influenza Data database (GISAID; www.gisaid.org) [3], representing 13% of all SARS-CoV-2 cases diagnosed in NSW at the time. The initiative to promptly release genomic data has mirrored other national and international efforts focused on near real-time monitoring of the evolution and intercontinental spread of the SARS-CoV-2 [4–6]. Prospective WGS of SARS-CoV-2 cases in NSW has continued and to date (April 30, 2021), 1710 genomes representing 32% of confirmed cases have been generated.

A large array of SARS-CoV-2 sequencing protocols have been developed since the start of the pandemic. They range from numerous target enrichment techniques which can be applied to multiple sequencing technologies, to suites of bioinformatics and data visualisation workflows [7]. The rapid development of all aspects of SARS-CoV-2 WGS was aided in part by efforts from the global genomics community in developing viral WGS methods (https://artic.network/ncov-2019). However, accurate SARS-CoV-2 genomic surveillance has been hampered by several common challenges: firstly, a high level of variability exists between sequencing protocols in obtaining complete SARS-CoV-2 genomes, particularly from clinical samples with low viral loads (as reflected by real-time PCR (RT-PCR) cycle threshold (Ct) values), such as those collected from patients without symptoms, mild disease, or late in the course of infection. Secondly, the accuracy required to detect and call variants using different protocols has not been adequately validated. All of these factors, sequencing method, reproducibility and thresholds for variant calling may affect the quality and impact of genomic surveillance and ultimately public health efforts to contain outbreaks.

Synthesis of SARS-CoV-2 genomic data with detailed epidemiological exposure and contact tracing information can provide definitive evidence of importation events and identification of local SARS-CoV-2 transmission chains [3, 8]. SARS-CoV-2 clusters, transmission chains, or networks linked to superspreading events are often differentiated genomically by single nucleotide polymorphisms (SNPs) within the SARS-CoV-2 genome [9]. The ability to rapidly and accurately characterise SNPs and other variants has become even more important after the identification of several ‘variants of concern’ (VOC). VOC contain specific mutations identified as important and relevant for COVID-19 control due to mounting evidence of positive selection of specific non-synonymous spike protein mutations that can increase the duration, severity and transmission of COVID-19 by affecting host immune responses [10–13]. Complete genomes generated using highly sensitive and specific sequencing methods are therefore required to inform and enable genomics guided surveillance to provide the information necessary for COVID-19 control and policy decisions, particularly as widespread SARS-CoV-2 vaccination is underway [14–17].

This study systematically evaluated three different sequencing methods for their sensitivity and ability to generate complete SARS-CoV-2 genome sequences suitable for public health surveillance. We assessed and compared (i) the pooled long-amplicon method [2] with (ii) ARTIC v3 network tiled amplicon protocol (https://artic.network/ncov-2019) that has been adopted widely since the start of the pandemic, and (iii) a probe capture-based Respiratory Viral Oligo Panel (RVOP) (Illumina). Additionally, we investigated the pattern of low frequency variants generated by these methods, which can be important in defining and highlighting transmission chains [18].

## RESULTS

### Viral isolates, viral loads and genome profiles

Seven SARS-CoV-2 positive clinical specimens were cultured as representatives of different SARS-CoV-2 genomic clusters that were co-circulating in NSW between February - April 2020 [3]. Details of the genome obtained from each clinical specimen, including GISAID ID, lineage and SNP profile are listed in supplementary Table 1. Two of seven isolates lost a SNP compared to the genome obtained directly from the original clinical specimens. The genome of Isolate 2 reverted to wild-type at position C:26213; however, the SNP C:26213:T detected in the original clinical specimen was still present as a low frequency variant. In Isolate 7, all reads at position 13730 were the wild-type allele (C). To investigate the effect of low viral load on detection of variants, serial dilutions of cell culture supernatant were performed. RT-PCR results from each culture dilution demonstrated that a 10-fold decrease in viral load corresponded to a Ct increase of ~3-4 cycles (Fig.1). A total of seven dilutions were made, five of which remained consistently SARS-CoV-2 RT-PCR positive with corresponding viral loads decreasing from a median load of 71,062 copies/μL (median Ct 25.42, range Ct 24.29 – 26.65, viral load range 47,482 – 1,178,540 copies/μL) to a median of 112 copies/μL (median Ct 36.62, range 34.7 – 38.19, viral load range 18 – 1,584 copies/μL). Culture dilutions with Ct >39 were deemed too low to attempt sequencing and were excluded from further analysis.

**Figure 1:**
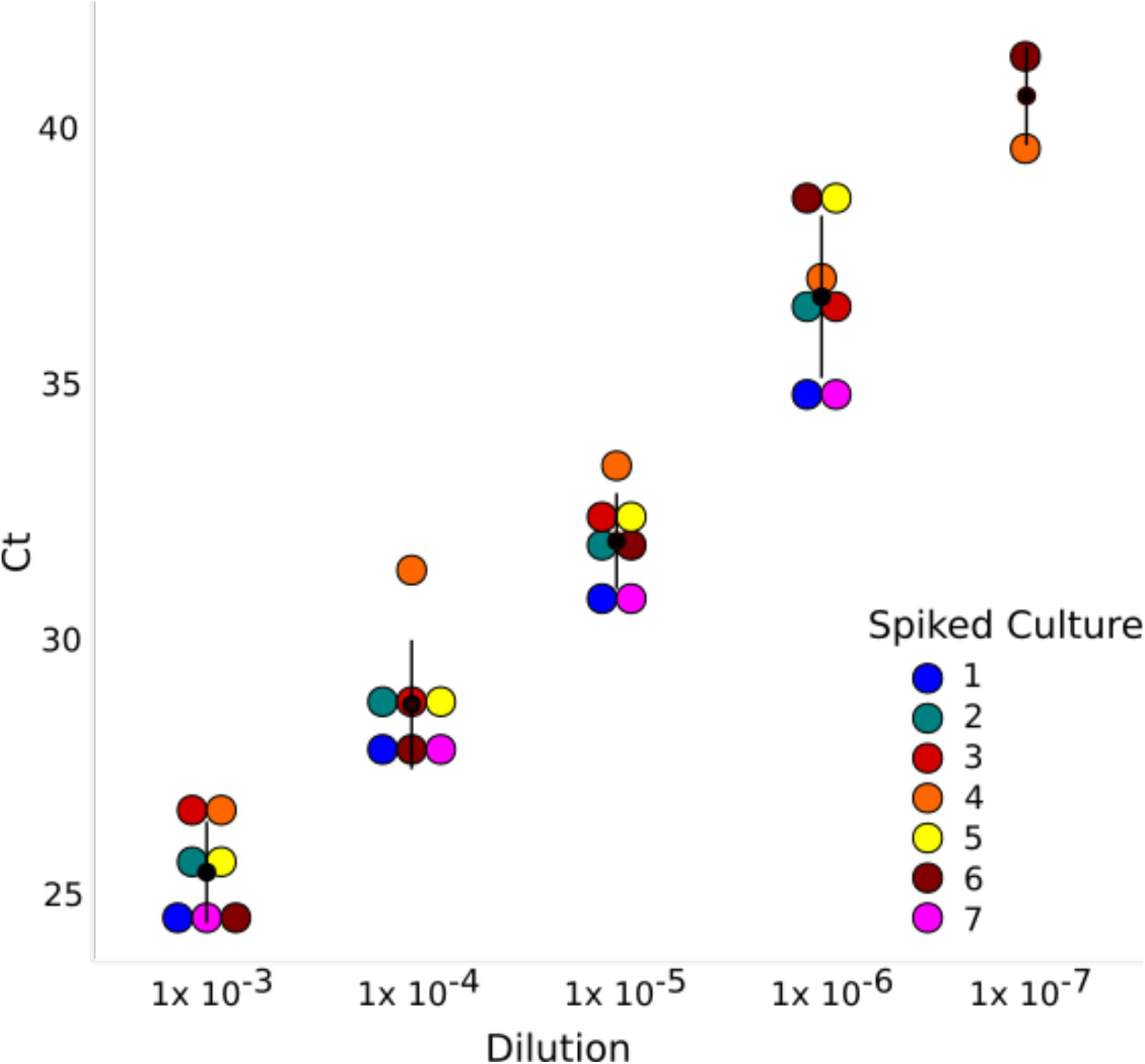
Viral load of SARS-CoV-2 cultures spiked in respiratory matrix. RT-PCR quantification of seven serially diluted SARS-CoV-2 cultures demonstrates an increase of 3-4 cycles for each ten-fold dilution of viral culture.

### Synthetic control

Using the long-amp method, only 57% (8/14 amplicons) of the synthetic control genome were able to be sequenced up to ~Ct 32, after which no amplicons were produced. Regions which did not amplify at higher viral loads were A2, A3, A4, B4, B5 and B6, signalling that that these primer pairs span two contiguous but separate segments of the synthetic genome. The smaller tiled amplicons from ARTIC v3 produced a higher proportion (93.9%, 92/98 amplicons) of the genome; however, amplicons 16, 17, 33, 50, 66 and 82 did not amplify. Missing regions from both ARTIC v3 and long-amplification methods overlapped, confirming six distinct segments of the synthetic control. Due to the non-amplification of larger products from the long-amplification method, less of the genome was able to be recovered, meaning that subsequent variant calling from these missing regions could not be performed. Complete genomes (>99% coverage) for the synthetic control was able to be obtained using RVOP up to Ct 28.

### Comparison of genome coverage across three sequencing methods

At Cts 25-29 (up to 2000 copies/μL), all three WGS methods generated near complete SARS-CoV-2 genomes with >10x coverage (Figure 2, Supplementary Figure 1). The highest level of genome coverage across all five dilutions was achieved using ARTIC v3 with >90% genome coverage achieved at viral loads down to ~Ct 38 (2 copies/μL). For each of the complete genomes (expected genome size of 29,903bp), there were fewer than 1000 ambiguous bases (N’s) from the MN90894.3 reference genome (Figure 2). On the other hand, genome coverage decreased substantially using long-amplicon and RVOP methods at a median Ct 32 (Ct range 30.7 – 33.4, median viral load 1340 copies/μL, range 725 – 14,613 copies/μL) (Supplementary Figure 1) however the differences observed were not significant (Figure 2). This decreasing trend continued at lower dilutions for both long-amp and RVOP, resulting in significant differences of the genome coverage obtained using the ARTIC, long-amplification and RVOP methods (p<0.05) (Figure 2).

**Figure 2.**
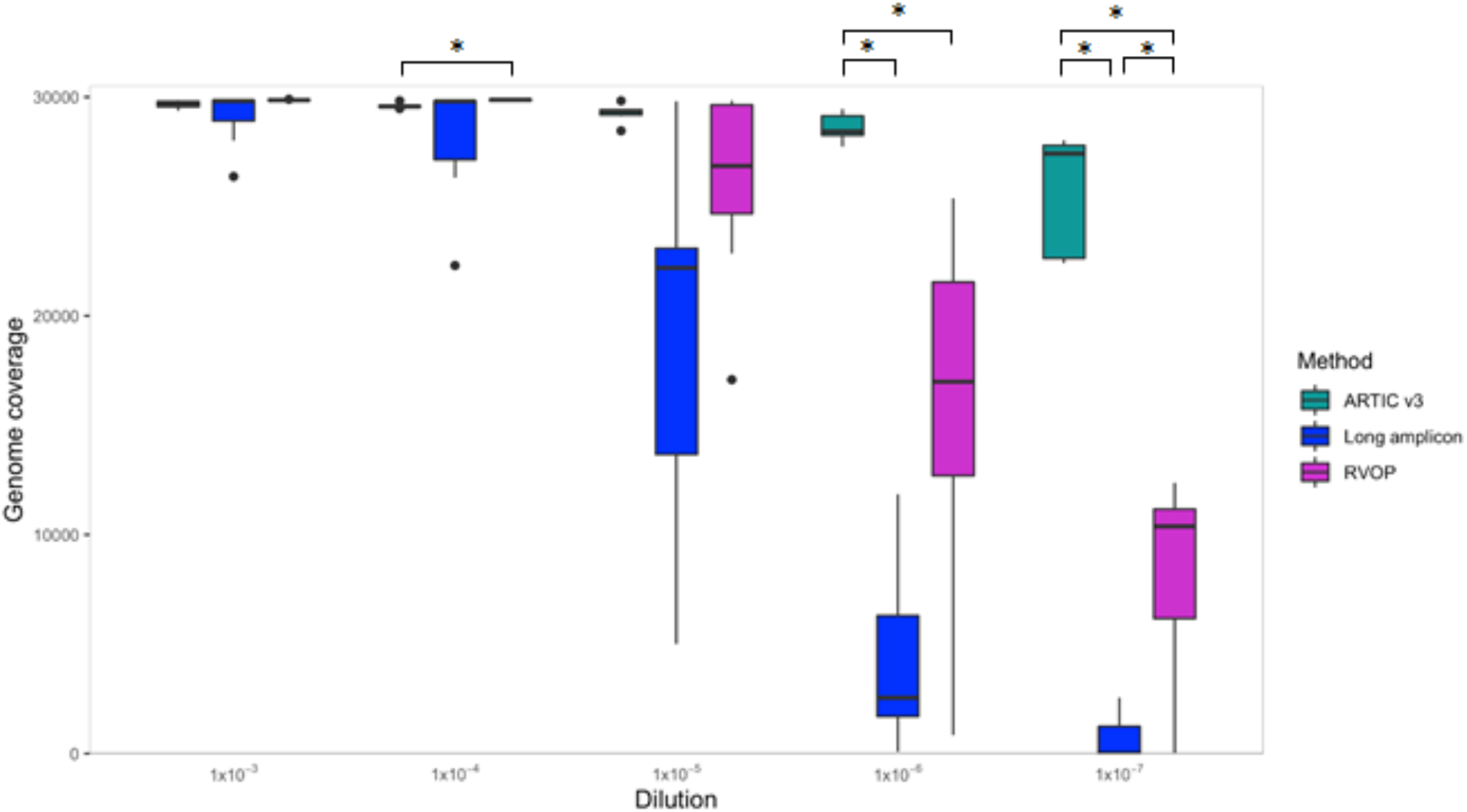
Boxplot showing the genome coverage of ARTIC v3, long-amplification and probe capture (RVOP) based whole genome sequencing methods of SARS-CoV-2 cultures. Significant differences (*) observed in genome coverage between different methods (p<0.05). NB pairwise comparisons between each methodology were only performed within each dilution.

### Read depth affects genome coverage and variant calling

Read depth across amplicons differed substantially between ARTIC v3 and long-amp methods creating highly uneven genome coverage. ARTIC v3 amplicons 9, 17, 23, 64, 67, 70, 74 and 91 were amplified inconsistently at higher Ct values (Ct >34). A2, B3 and B6 from the long-amplification protocol were the poorest performing, often not amplified in samples with Ct <30. These 400 bp-5 kb missing amplicons created large genomic gaps, which made variant calling problematic. In contrast, the amplicons which amplified with high efficiency using ARTIC v3 (44, 57, 62), had consistently higher average read depths regardless of Ct value. The RVOP achieved the most consistent read depth across the genome, with relatively even distribution of missing bases compared with either amplification-based sequencing method. However, average read depth of samples at ~Ct 32 (Ct range 30.7 – 33.4) was low (Figure 3) with inconsistent genome coverage <10x, also resulting in problems with variant calling.

**Figure 3:**
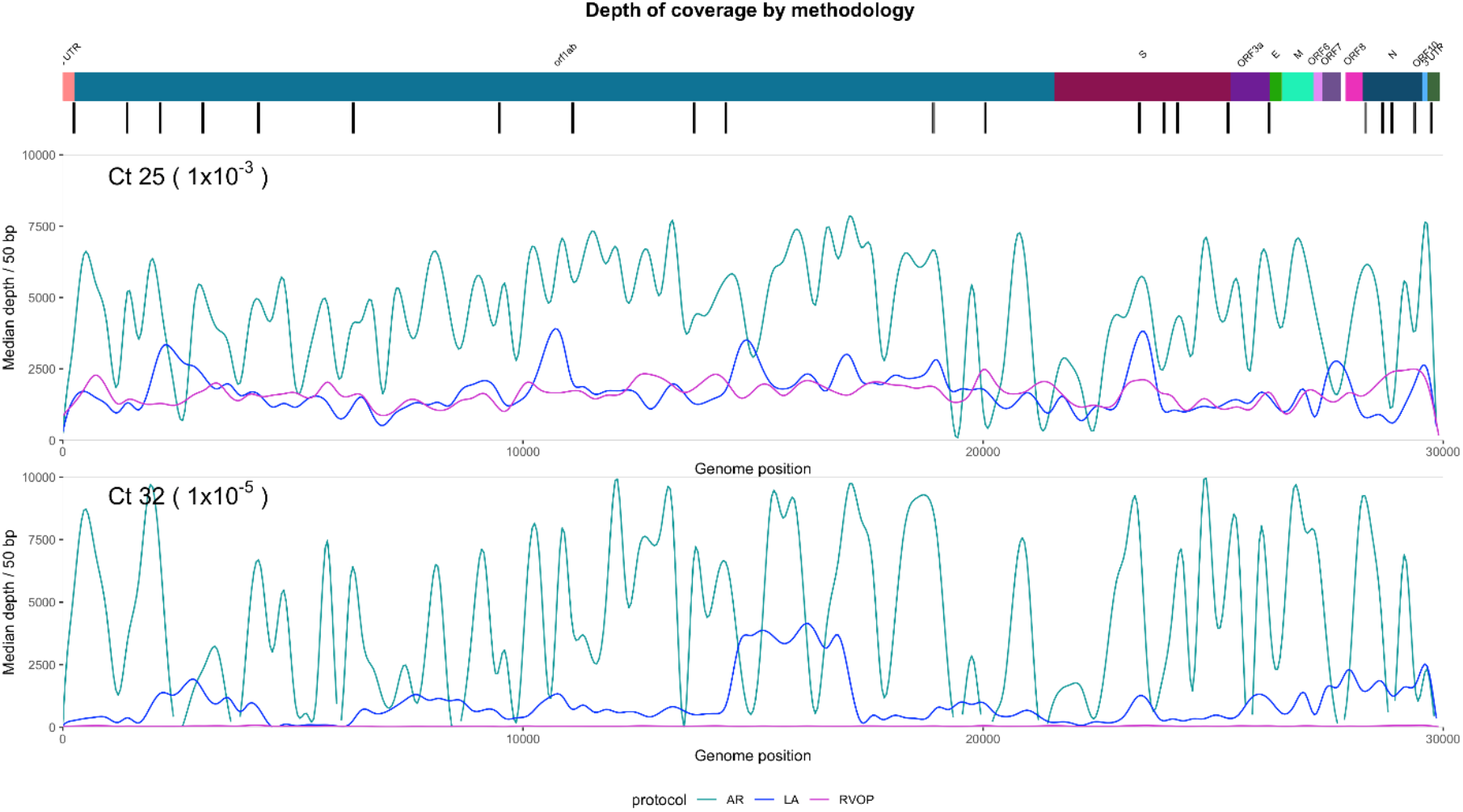
Overall read depth across the SARS-CoV-2 genome using ARTIC v3 (green line), long-amplification (blue line), and probe capture RVOP (pink line) whole genome sequencing methods. Depth is averaged across all samples for each method separately. Lines are smoothed by using the ‘geom_spline’ function in R. The coloured bar at the top represents the regions of the SARS-CoV-2 genome, and black bars represent informative single nucleotide polymorphisms.

### ARTIC rebalanced pools

Using the COVID-19 Genomics Consortium (COG-UK) guidelines, we rebalanced ARTIC v3 primers in an attempt to improve amplification of specific amplicons and obtain more even sequencing coverage across the genome. Figure 4 shows the performance of rebalanced primers compared with original primer concentrations prior to rebalancing. Unsurprisingly, as viral load decreases, coverage across poorer performing amplicons decreased in parallel (Figure 4 and Supplemental Figure 1). No significant changes in coverage were observed (across all dilutions) with amplicons 15, 27, 73, even though the primer concentrations were increased by 1.5x-2.1x. However, amplicons 64, 67, 70 and 74 (for which primer concentrations were increased by a factor of between 6-7.8x), performed significantly better than original unbalanced primer pools. Other amplicons (i.e. 36, 54 and 66) whose primers were increased by a factor of >3x performed worse than expected (likely due to potentially poorly designed primers or excessive interactions with other primers). Regardless of individual primer rebalancing factors, sufficient depth (>10x) to meet variant calling QC at Ct 35 was obtained for all amplicons.

**Figure 4:**
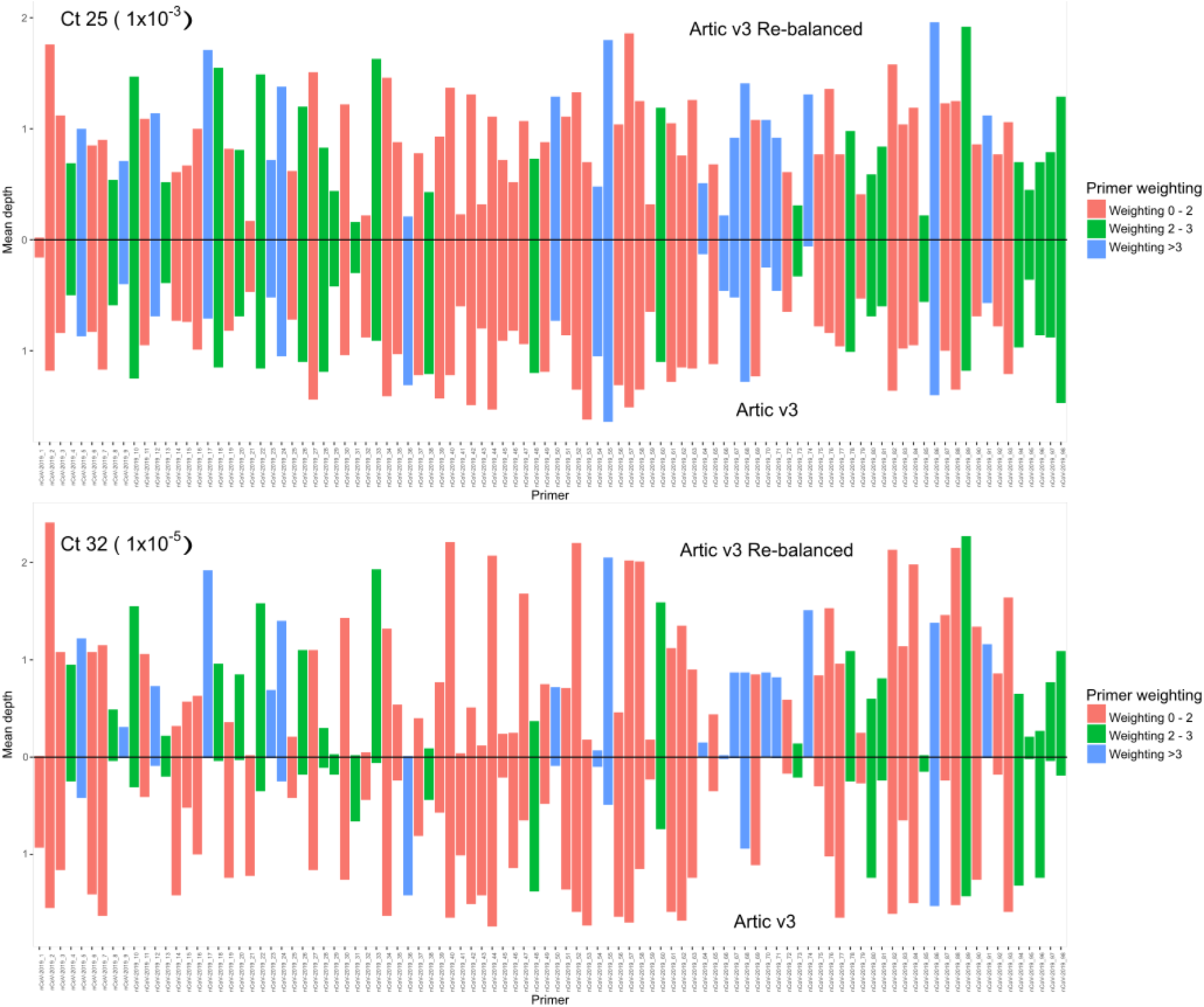
Comparison between ARTIC v3 original primer pooling and rebalanced primer pools at Ct 25 and Ct 32. Median depth is expressed as a factor of average read depth across each amplicon. Bars above zero represent sequencing depth achieved by ARTIC v3 rebalanced primer pools; height of bars below zero represent sequencing depth of ARTIC v3 original primer pools. ARTIC v3 primers are listed across the x-axis in sequential order across the genome. For exact primer weightings, refer to: https://www.protocols.io/view/covid-19-artic-v3-illumina-library-construction-an-bky5kxy6).

### Comparative sensitivity of three SARS-CoV2 sequencing methods

Sensitivity of each method was defined as the ability to accurately call SNPs, based on a clear consensus amongst all the dilutions. All three methods exceeded 90% sensitivity with median Ct 28.7 (Ct range 27.6 - 31.3, median viral load 12,025 copies/μL) (Figure 5). The sensitivity for ARTIC remained high for samples up to Cts >38, whereas sensitivities for both pooled long-amplicon and RVOP dropped below 80% at Ct >30. No differences in sensitivity or specificity were observed between ARTIC original primer pooling compared with rebalanced primer pools. Interestingly, at higher Ct values, the RVOP was more likely to represent true SNPs as low frequency variants (discussed in further detail below).

**Figure 5:**
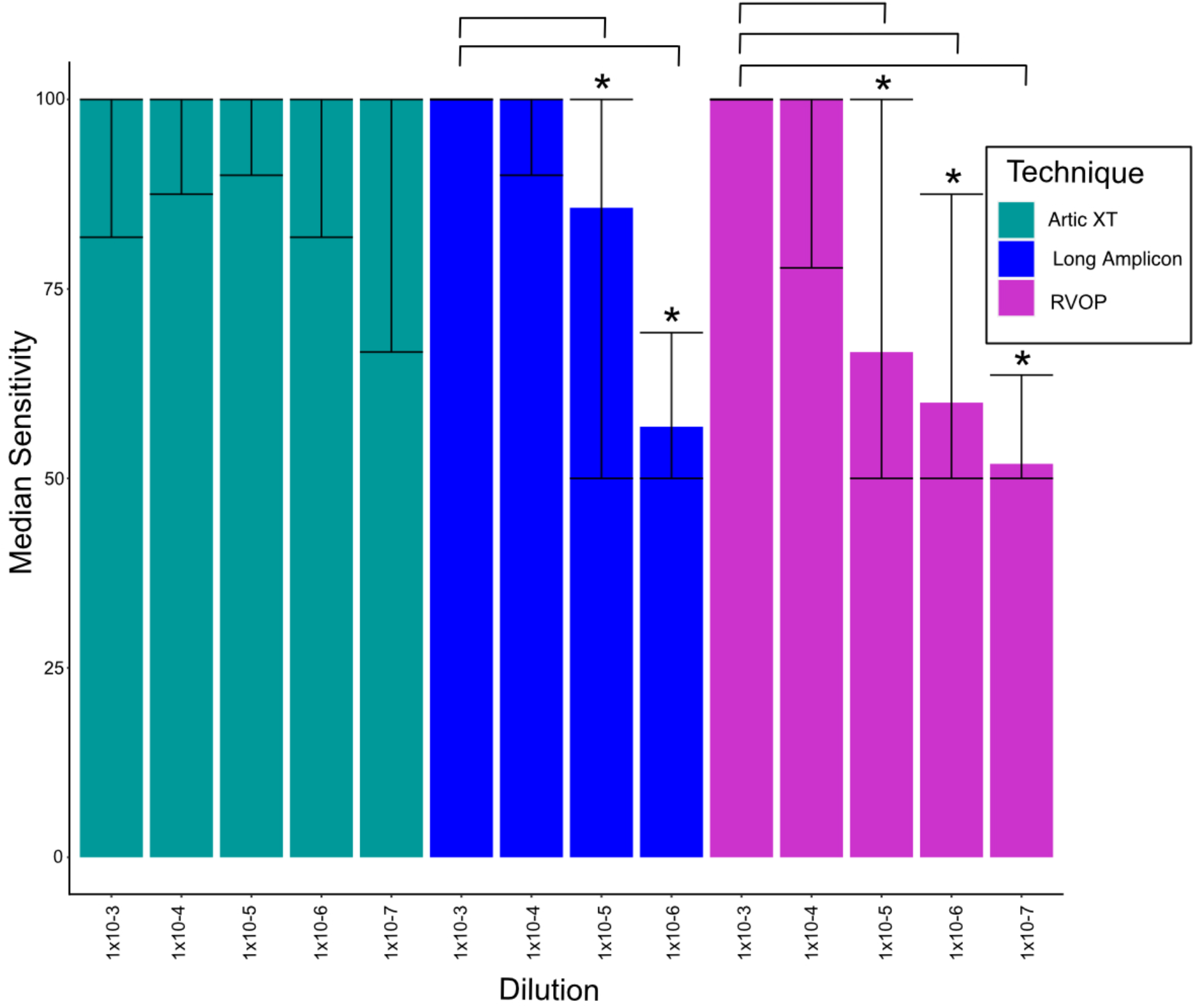
Sensitivity of ARTIC v3, long-amplification and RVOP (Respiratory Viral Oligo Panel, Illumina) whole genome sequencing methods. The sensitivity of ARTIC v3 was the highest across all viral load dilutions. No sensitivity calculations could be made for the pooled long-amplification method at 10^−7^ dilutions due to insufficient amplicons for variant calling. Significant differences (*) in sensitivity were observed between dilution 10^−3^ and dilutions 10^−5^ and 10^−6^ using the long-amplification method and dilution 10^−3^ and dilutions 10^−5^, 10^−6^ and 10^−7^ using the RVOP method. No differences were observed between ARTIC v3 original primer pooling and rebalanced pools.

### RVOP and the detection of other respiratory pathogens

The RVOP can detect 43 common human respiratory viruses and 60 human control genes (which serve as internal positive controls for library construction and sequencing steps) in individual clinical samples [19]. Trimmed reads from all seven diluted cultures prepared using the RVOP were mapped against 203 reference sequences of 43 respiratory pathogens. Human rhinovirus 89 (NC_001617.1) and adenovirus C (NC_001490.1) were detected, although coverage across both viral genomes was less than 2%. An in-house respiratory panel RT-PCR reconfirmed the presence of both rhinovirus (Ct 27) and adenovirus (Ct 26) in the negative respiratory matrix (supplementary Table 3). Twenty-seven reads mapped to human coronavirus 229E, but when BLAST was used to check the identity of these reads, the majority of mis-mapped reads also had high homology with SARS-CoV-2 and were subsequently found to have short read lengths (<40 bp).

### Low frequency variant detection

On average 16.7 low frequency variants were detected using all three techniques per sample (range 12-25). However, almost half of these low frequency variants were removed, due to their detection in a single dilution per isolate. Generally, these non-replicated low frequency variants were only detected in low viral load dilutions (1× 10-6 and 1×10-7). Low frequency variants repeatedly detected in at least two dilutions were most commonly detected using RVOP (median number of sites 10, range 6-16) followed by ARTIC (median number of sites 1, range 0-5) and long-amplification (median number of sites 1, range 0-4) and (Supplementary Figure 1, supplementary Table 2). The presence of low frequency variants was confirmed, at the same genome position by all three methods, in two culture isolates (median number of sites 2, range 0-4): Isolate 1 at positions 657, 27972, 29585, Isolate 2 at positions 12299, 16466 (supplementary Table 2). No additional low frequency variants were detected using the ARTIC v3 rebalanced pools. Despite using a simulated respiratory matrix to control for background artefacts, there was little consistency in the number and location of low-frequency variants detected across the diluted genomes using each of the three methods.

## DISCUSSION

This study highlighted important quality requirements for high-throughput sequencing of SARS-CoV-2 for the purpose of public health surveillance. These parameters are critical for the application of SARS-CoV-2 genomics in tracking transmission pathways and monitoring ongoing viral evolution in circulating virus populations. Sequencing of samples with low viral loads and high Ct values (eg. Ct >33) has been challenging regardless of methodology used [20–23]. Sequencing of such samples can still be attempted, but the resulting genomes often have a substantial portion of missing bases, making it difficult to infer genomic clusters or identify VOC.

Our findings demonstrated the rapid loss of genome coverage using pooled long-amplicon sequencing and the RVOP at Ct >32 (median viral load 1340 copies/μL), indicating that low viral load or suboptimal RNA quality can be a limiting factor that must be considered when using these methods to generate reproducible genomic data. In contrast, near complete genomes can be recovered using ARTIC v3 at Ct >38, suggesting that the ARTIC protocol is either more sensitive at low viral loads or less impacted by reduced RNA quality. Indeed, the ARTIC protocol has performed well for samples with higher viral loads (Ct <25) [24–26] and has been implemented in numerous laboratories worldwide. However, at lower viral loads, we found both amplification-based methods inconsistently produced data in genomic regions of known significance. Analogous to the findings presented here, uneven amplification efficiencies and coverage bias have been widely reported for low viral load specimens [25, 26]. Increasing coverage over underperforming regions of the genome may be achieved by sequencing at higher depths, but this approach is costly and impractical in outbreak situations where high and rapid throughput is necessary. Rebalancing primer concentrations for ARTIC v3 improved coverage over previously poorly sequenced regions, and it is likely that additional manipulation of primer pooling or primer design would further enhance coverage.

In contrast, the RVOP generates consistent and even SARS-CoV-2 genome coverage over a range of Ct values, despite the sensitivity being only marginally higher than for long-amplicon sequencing. While not examined fully in this study, the RVOP can simultaneously detect other pathogens in a single sample, reducing delays in diagnosis and treatment options for patients who test negative for SARS-CoV-2. Similar to the genome coverage achieved for SARS-CoV-2 in this study, the RVOP should also be able to generate whole genomes of other respiratory viral pathogens targeted by the panel. We were unable to confirm complete coverage of adenovirus and rhinovirus (despite their presence confirmed using RT-PCR) as the pooled respiratory matrix used for this study consisted of a convenient sample of SARS-CoV-2 negative universal transport media (UTM). Poor sample quality as a result of suboptimal transport and storage conditions may have been another contributing factor to the limited and inconsistent coverage of other respiratory pathogens.

The loss of informative sequencing data, especially in genomic regions of interest, can hamper public health efforts to monitor changes in circulating viral populations. Given that numerous VOC have been identified worldwide [14, 27–29], amplicon drop-outs, particularly within the spike region, are problematic. For instance, B6 from the long-amplification protocol, and amplicons 70 and 74 from the original ARTIC v3 protocol encompass part of the spike protein, but all performed poorly and often did not amplify at Ct >32. Rebalancing the ARTIC v3 primer pools increased sequencing coverage and depth over amplicons 70 and 74. However, it is important to note that both long-amplicon and ARTIC v3 methods involve primer binding prior to amplification and are therefore prone to amplicon drop-outs if variants arise within primer sites. The risk of amplicon dropouts can be overcome by redesigning primers away from variant sites; such protocol changes can be time consuming and difficult to implement but will be necessary given the rapid rise and spread of VOC. The constantly changing population dynamics of the circulating SARS-CoV-2 viruses will require ongoing, high quality genomic surveillance to track the evolution of circulating isolates and help inform necessary changes to sequencing methodologies.

Detecting and locating genomic positions of low frequency variants from culture derived specimens can provide insight into the reliability of intra-host single nucleotide variants (iSNVs) called from clinical specimens. The role of intra-host genomic variability in SARS-CoV-2 may be important in inferring transmission events [30] and may be responsible for significant complications in patients with malignancies [31]. Thus, such low frequency variants require ongoing detection and surveillance. There have been suggestions that iSNVs can be detected at a frequency as low as 2% [18], however only those iSNVs occurring at a frequency of >10% and a minimum coverage of 100x were investigated in the present study. At this threshold, substantial variability of low frequency variants was observed using the methods tested in this study even after controlling for background artefacts generated during the WGS process (via the use of viral cultures in a defined respiratory matrix). The inconsistency in low frequency variants calls can be attributed to the unique sequencing chemistries of each method, and the impact of upstream amplification and hybridisation procedures, highlighting the importance of recognising and accounting for biases that arise during both laboratory preparation and downstream bioinformatic processes.

While we have systematically tested and determined the threshold at which complete genomes can be generated for each method, we have not yet addressed issues with poor quality specimens. Quality and quantities of RNA in clinical specimens for WGS is highly dependent on sample types, collection methods, transport and processing. Suboptimal processes are not uncommon and inherent with high throughput and often centralised testing. Sample degradation as a result of these factors has been highlighted as a significant problem to generating high quality genome sequences [23].

In conclusion, our systematic evaluation of sensitivity and ability to detect low frequency variants demonstrated that overall, the ARTIC v3 protocol was the most sensitive method for generating complete SARS-CoV-2 genomes. The additional advantages of the ARTIC protocol are better capacity to recover genomes from clinical samples with low viral loads and the ability to detect low frequency variants. Ongoing updates to the ARTIC v3 protocol, such as the rebalancing of primer pools (through the COG-UK and efforts from research institutions), will ensure continual improvements to the WGS process. The optimisation of SARS-CoV-2 genome sequencing can increase the utility of SARS-CoV-2 genomics for COVID-19 cluster detection, transmission tracking and public health responses.

## METHODS

### Clinical specimens

The study period and region included the 4 months between March and July 2020, in NSW, Australia. SARS-CoV-2 RT-PCR positive specimens which were subsequently cultured at NSW Health Pathology-Institute of Clinical Pathology and Medical Research (ICPMR) in the study period were included for selection. Respiratory samples in UTM which were RT-PCR negative for SARS-CoV-2 were collected and stored at 4°C. These negative specimens were de-identified and pooled together totalling 40 mL before RNA was extracted. This RNA was used to dilute SARS-CoV-2 isolates, referenced in the manuscript as negative respiratory matrix. Ethical and governance approval for the study was granted by the Western Sydney Local Health District Human Research Ethics Committee (2020/ETH02426).

### Viral isolation

SARS-CoV-2 positive respiratory specimens were cultured in Vero C1008 cells (Vero 76, clone E6, Vero E6 [ECACC 85020206]) as previously outlined [32]. Briefly, Vero cell cultures were seeded at 1-3 × 10^4^ cells/cm^2^ in Dulbecco’s minimal essential medium (DMEM, LONZA, Alpharetta, GA, USA) supplemented with 9% foetal bovine serum (FBS, HyClone, Cytiva, Sydney, Australia). Media was replaced within 12 hours with inoculation media containing 1% FBS with the addition of penicillin, streptomycin and amphotericin B deoxycholate to prevent microbial overgrowth and then inoculated with 500 μL of SARS-CoV-2 positive respiratory sample. The inoculated cultures were incubated at 37°C in 5% CO_2_ for 5 days (days 0 to 4). Cell cultures were observed daily for cytopathic effect (CPE). Routine mycoplasma testing was performed to exclude mycoplasma contamination of the cell line and all culture work was undertaken in physical containment laboratory level 3 (PC3) biosafety conditions. The presence of CPE and increasing viral load was indicative of positive SARS-CoV-2 isolation. RT-PCR testing was performed on day 1, 2, 3 and 4 by conducting RNA extraction and SARS-CoV-2 RT-PCR on 200 μL of culture supernatant. Culture supernatant was harvested four days after inoculation and stored at −80°C.

### RNA extraction from viral culture

A total of 600 μL (three × 200 μL) of day 4 SARS-CoV-2 culture supernatant was used as input into the RNeasy Mini Kit (Qiagen) for RNA extraction with minor modifications. 600 μL of RLT buffer was added to 200 μL of sample and mixed well. An equal volume (800 μL) of 70% ethanol was then added and mixed well by pipetting, before loading onto RNeasy column in successive aliquots until the entire volume was extracted. RNA was eluted in 30 μL, pooled for a total of 90 μL and stored at −80°C prior to dilution. Total RNA was extracted from pooled SARS-CoV-2 negative clinical specimens as above.

### Respiratory virus detection by RT-PCR

A previously described RT-PCR [33] targeting the N gene was employed to estimate the viral load of cultured RNA and ensure the absence of SARS-CoV-2 in the negative respiratory matrix. Additional RT-PCRs were used to investigate the presence of common viral respiratory viruses: human influenza viruses A & B, parainfluenzaviruses 1, 2 & 3, respiratory syncytial virus, adenovirus, and rhinovirus in negative UTM extract [34].

### Synthetic control

A commercially available synthetic RNA control reference strain (Wuhan - 1 strain, TWIST Biosciences) containing six non-overlapping fragments replicating the most commonly used reference sequence (NCBI GenBank accession MN908947.3) was used as a control. Serial 10-fold dilutions starting at 20,000 copies/μL to 2 copies/μL were made and used to generate a standard curve and quantify the viral load of each culture spiked dilution per reaction. The N gene SARS-CoV-2 RT-PCR was used to determine the viral load of the neat culture RNA after extraction. The synthetic control was also serially diluted 10-fold in respiratory matrix (as outlined below), enriched using each of the methods below and sequenced in parallel with diluted cultures.

### Normalisation and serial dilution of viral culture RNA into negative respiratory matrix

Based on the viral load of the neat culture RNA (Ct 12.57– 14.48, viral load 2.0×10^8^ - 6.0×10^7^ copies/μL), each culture RNA extract was diluted 1:10 with negative RNA extract. Then 10-fold serial dilutions were made in negative RNA extract until an estimated concentration of >10 copies/μL (Ct 37-40) was reached for each isolate. cDNA was generated for all serially diluted RNA samples using LunaScript RT SuperMix Kit (New England BioLabs). Sufficient volume was prepared to perform duplicates for each method at each dilution. RNA and corresponding cDNA dilutions were aliquoted and stored at −80°C and −20°C, respectively. RT-PCR was then performed for each sample dilution to determine Ct value and corresponding viral load.

### Viral enrichment and genome sequencing

For each of the serially diluted samples, viral enrichment was performed using three methods: ARTIC v3, a 14-pool long-amplicon approach, and probe capture using Illumina RNA Prep with Enrichment with the Respiratory Viral Oligo Panel (RVOP). Details of each enrichment method are outlined below.

#### ARTIC v3 nCoV-2019 sequencing protocol - ARTICv3 (https://www.protocols.io/view/ncov-2019-sequencing-protocol-v3-locost-bh42j8ye)

The ARTIC v3 protocol was performed with the following modifications. Tiling PCR was used to amplify the whole genome according to ARTIC nCoV2-2019 sequencing protocol. Each PCR included 12.5 μL Q5 High Fidelity 2x Master Mix (New England Biolabs), 3.6 μL of either pool 1 or pool 2 10μM primer master mix (final concentration of each primer was ~10-11pM), 5 μL of template, molecular grade water was added to generate a total volume of 25 μL. Cycling conditions were as follows: initial denaturation at 95°C-2 min, followed by 35 cycles of: 95°C for 30 s, 63°C for 2 min 45 s, and a final extension step of 75°C for 10 min. Pool 1 and pool 2 amplicons were combined and purified with a 1:1 ratio of AMPureXP beads (Beckman Coulter) and eluted in 30 μL of sterile water. Purified products were quantified using Qubit™ 1x dsDNA HS Assay Kit (ThermoFisher Scientific) and diluted to the desired input concentration for library preparation. Sequencing libraries were prepared using Nextera XT (Illumina) according to manufacturer’s respective instructions. Sequencing libraries were then sequenced as 2×150bp runs on either the Illumina iSeq or MiniSeq platforms.

An updated ARTIC v3 protocol with rebalanced primer pools was also evaluated in this study. Primers for each ARTIC v3 pool were combined according to updated COG-UK consortium guidelines (https://www.protocols.io/view/covid-19-artic-v3-illumina-library-construction-an-bky5kxy6). Subsequent PCR and sequencing using the rebalanced ARTIC primer pools were performed as above.

#### Pooled long-amplicon PCR (dx.doi.org/10.17504/protocols.io.befyjbpw)

Pooled long-amplicon sequencing was performed as described previously [2]. Briefly, 14 overlapping PCR amplicons were independently generated and pooled together in equal volumes. Pooled products were purified with 0.8x AMpure XP beads (Beckman Coulter) and eluted in 30 μL of sterile water. Qubit™ 1x dsDNA HS Assay Kit (ThermoFisher Scientific) was used to quantify pooled amplicons before diluting to the desired input concentration for library preparation. Sequencing libraries were prepared using the Nextera XT kit (Illumina) and sequenced on either iSeq or MiniSeq (Illumina) using 2×76bp paired end reads. No other changes were made to the protocol.

#### Respiratory Viral Oligo Panel (RVOP)

Diluted culture RNA extracts were used as input into the RNA Prep with Enrichment kit (Illumina). RNA denaturation, first and second strand cDNA synthesis, cDNA tagmentation, library construction, clean up and normalisation were performed according to manufacturer’s instructions. Individual libraries were then combined in 3-plex reactions for probe hybridisation. The Respiratory Viral Oligo Panel v2 (Illumina) was used for probe hybridisation with the final hybridisation step held at 58°C overnight. Hybridised probes were then captured and washed according to manufacturer’s instructions and amplified as follows: initial denaturation 98°C for 30 s, 14 cycles of: 98°C for 10 s, 60°C for 30 s, 72°C for 30 s, and a final 72°C for 5 min. Library quantities and fragment size were determined using Qubit™ 1x dsDNA HS Assay and Agilent HS Tapestation and sequenced using 2×76bp runs on the Illumina MiniSeq.

### Bioinformatic analysis

Raw sequence data was processed using an in-house quality control procedure prior to further analysis. Demultiplexed reads were quality trimmed using Trimmomatic v0.36 (sliding window of 4, minimum read quality score of 20, leading/trailing quality of 5 and minimum length of 36 after trimming) [35]. Reference mapping and variant calling was performed using iVar version 1.2 [36]. Briefly, reads were mapped to the reference SARS-CoV-2 genome (NCBI GenBank accession MN908947.3) using BWA-mem version 0.7.17, with unmapped reads discarded. Primer positions were supplied to iVar trim to soft-clip any reads in the bam file which matched primer sequences. Average genome coverage was estimated by determining the number missing bases (N’s) in each sequenced genome. Variants were called using iVar variants (min. read depth >10x, quality >20, min. frequency threshold of 0.1). SNPs were defined based on an alternative frequency ≥0.9 whereas low frequency variants were defined by an alternative frequency between 0.1 and 0.9. Low frequency variants with <100x depth were excluded over concerns over reliability of calls where the frequency of either allele dropped below 10. Low frequency variants were only included if they were detected in 2 or more dilutions of each spike culture sequenced. Variants falling in the 5’ and 3’UTR regions were excluded due to poor sequencing quality of these regions. Polymorphic sites that have previously been highlighted as problematic were monitored [37]. SARS-CoV-2 lineages were inferred using Phylogenetic Assignment of Named Global Outbreak LINeages v2 (PANGOLIN) (https://github.com/hCoV-2019/pangolin) [38]. The frequency and positions of polymorphisms were compared between dilutions of the same culture and also against the original genome generated from the respiratory specimen and between cultures. Median genome coverage was calculated using the median depth in 50 bp bins across the reference genome for each method and dilution. Median read depth per amplicon was assessed in non-overlapping segments of each ARTIC v3 amplicon which was then converted to a factor of the expected read coverage (total mapped reads/genome size*150 bp). These factors were compared between original and rebalanced ARTIC v3 sequencing runs. Graphs were generated using R (version 3.6.1).

### Analytical performance-sensitivity and specificity

Sensitivity and specificity were calculated for each sequencing method using a consensus SNP approach. For each isolate, a SNP called in any method was considered a ‘true positive’ SNP if it occurred in two or more sequencing methods at the highest dilution. SNPs identified by a single sequencing method only (and not detected in the original clinical specimen) were considered false positives. Sensitivity was calculated using the following formula A/(A+C) × 100 where A was the number of true positive SNPs and C is the number of false negative SNPs. Specificity was calculated using the formula D/(D+B) x100 where D was the number of true negative bases (within the CDS region) and B was the number of false positive SNPs. Pairwise statistical comparisons were conducted between genome coverage and sensitivities at each dilution across each method using Friedman Test or Mann-Whitney tests with a significance level of p<0.05.

## Supporting information

SRA Accession details for fastQ files generated in this study

## Acknowledgements

The authors thank the Sydney Informatics Hub at The University of Sydney, NSW Ministry of Health, and staff from the Microbial Genomics Reference Laboratory and the Virology Laboratory, NSW Health Pathology-ICPMR for their assistance. We are grateful to partner laboratories within the NSW Health Pathology network and private laboratories for their participation in the COVID-19 genomics surveillance and to GISAID for collating international data on SARS-CoV-2.

## Data availability

Fastq files have been deposited in BioProject PRJNA723901 for all 118 genomes produced in this study. Individual SRA and GISAID accessions can be found in Supplementary Table 4 and 1 respectively.

---

**Supplementary materials**

**Supplementary Table 1:**
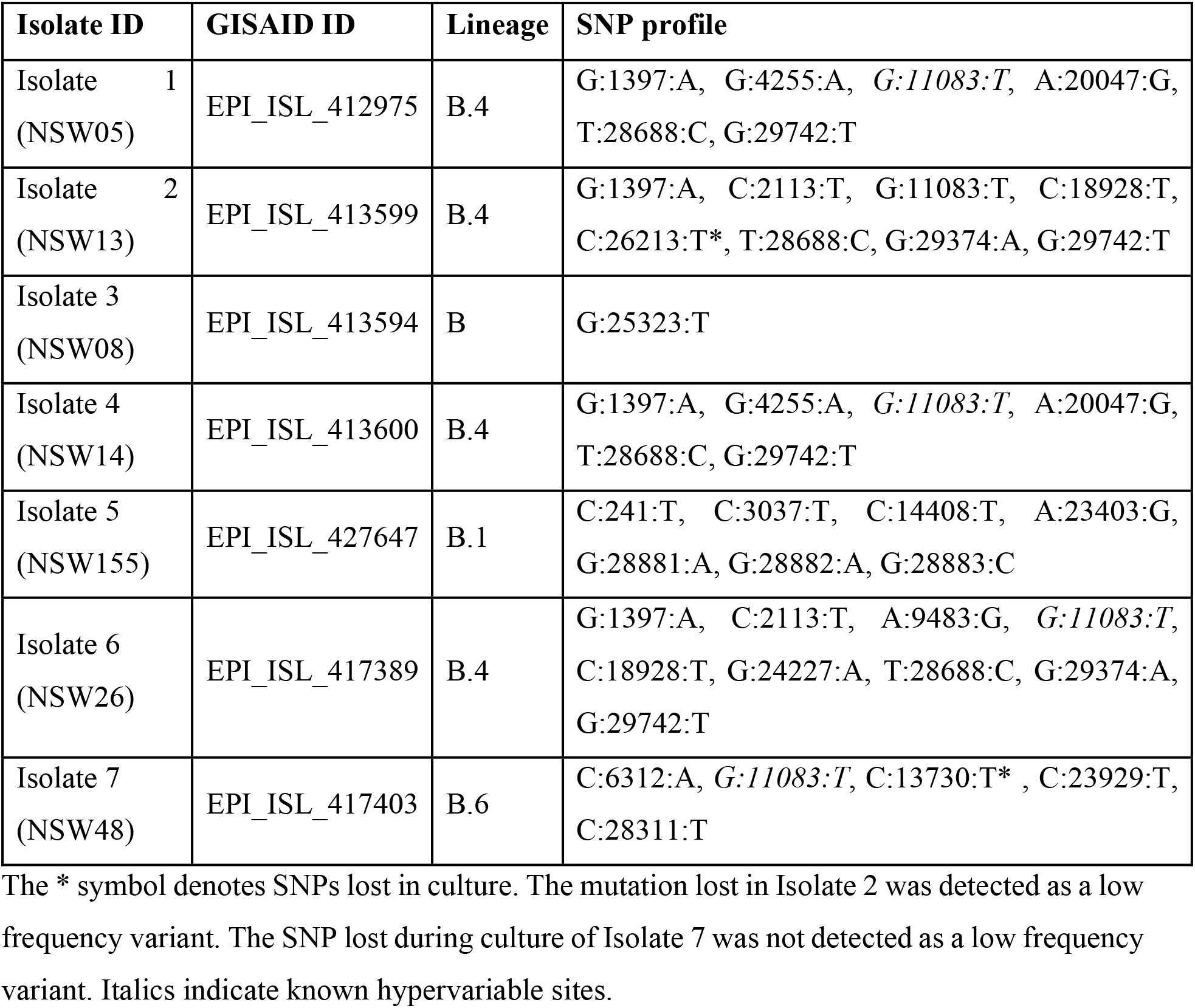
Genomes of SARS-CoV-2 isolates used in the study.

**Supplementary Table 2.**
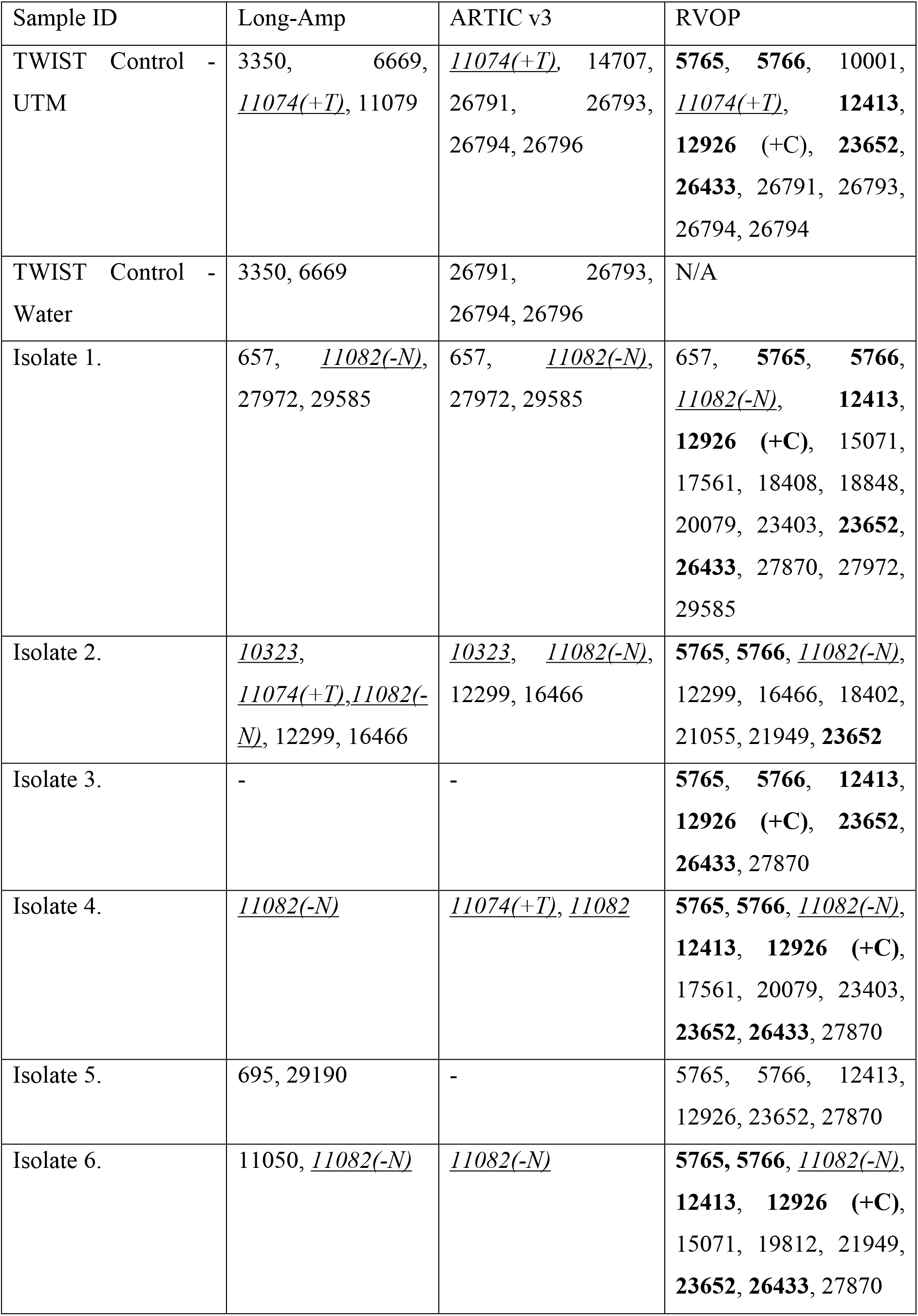
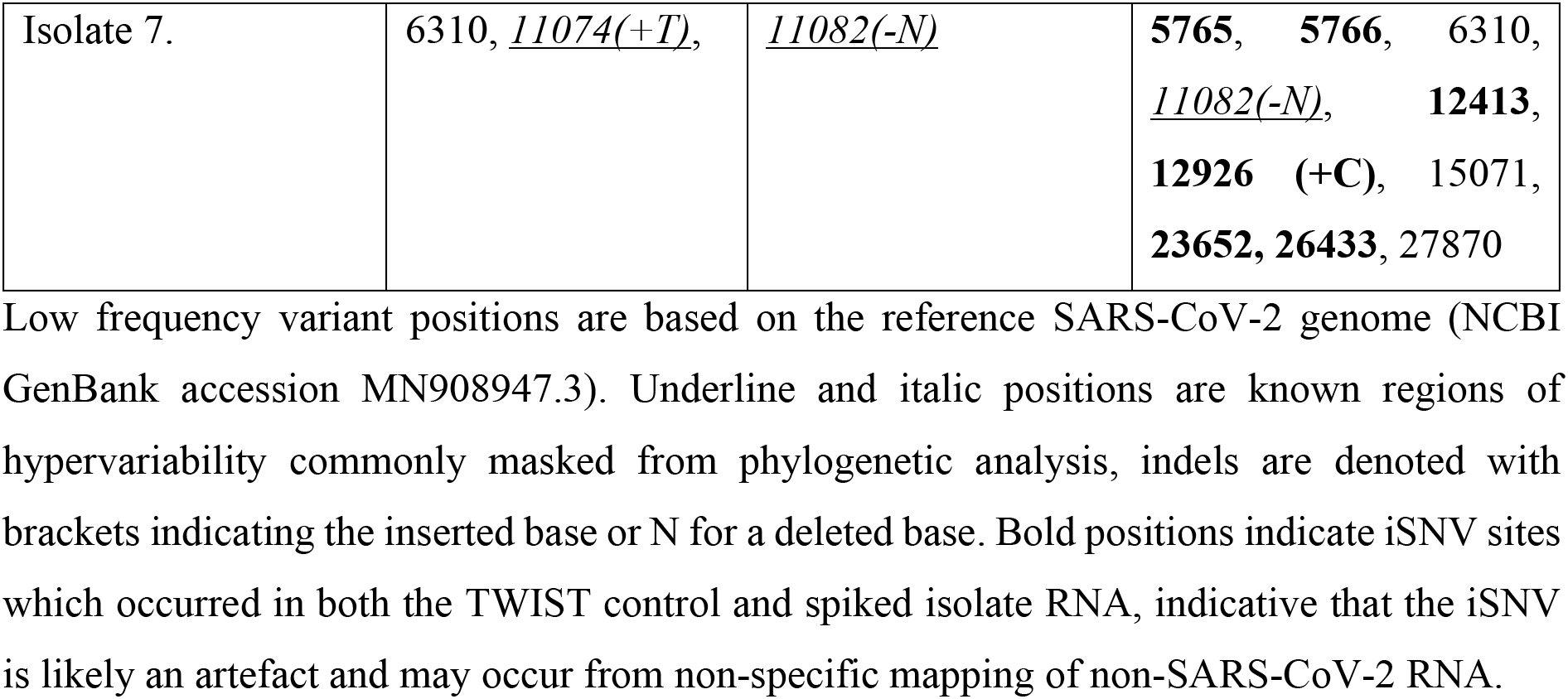
Low frequency variants detected using each method.

**Supplementary Table 3:** Negative SARS-CoV-2 RNA extract and detection of other respiratory pathogens

| Respiratory pathogen | Ct Value |
| --- | --- |
| Rhinovirus | 27 |
| Adenovirus | 27 |
| Influenza | ND |
| RSV | ND |
| Parainfluenza | ND |

**Supplementary Table 4: Details of SRA data availability for all 118 genomes produced in the study**

**Supplementary Figure 1:**
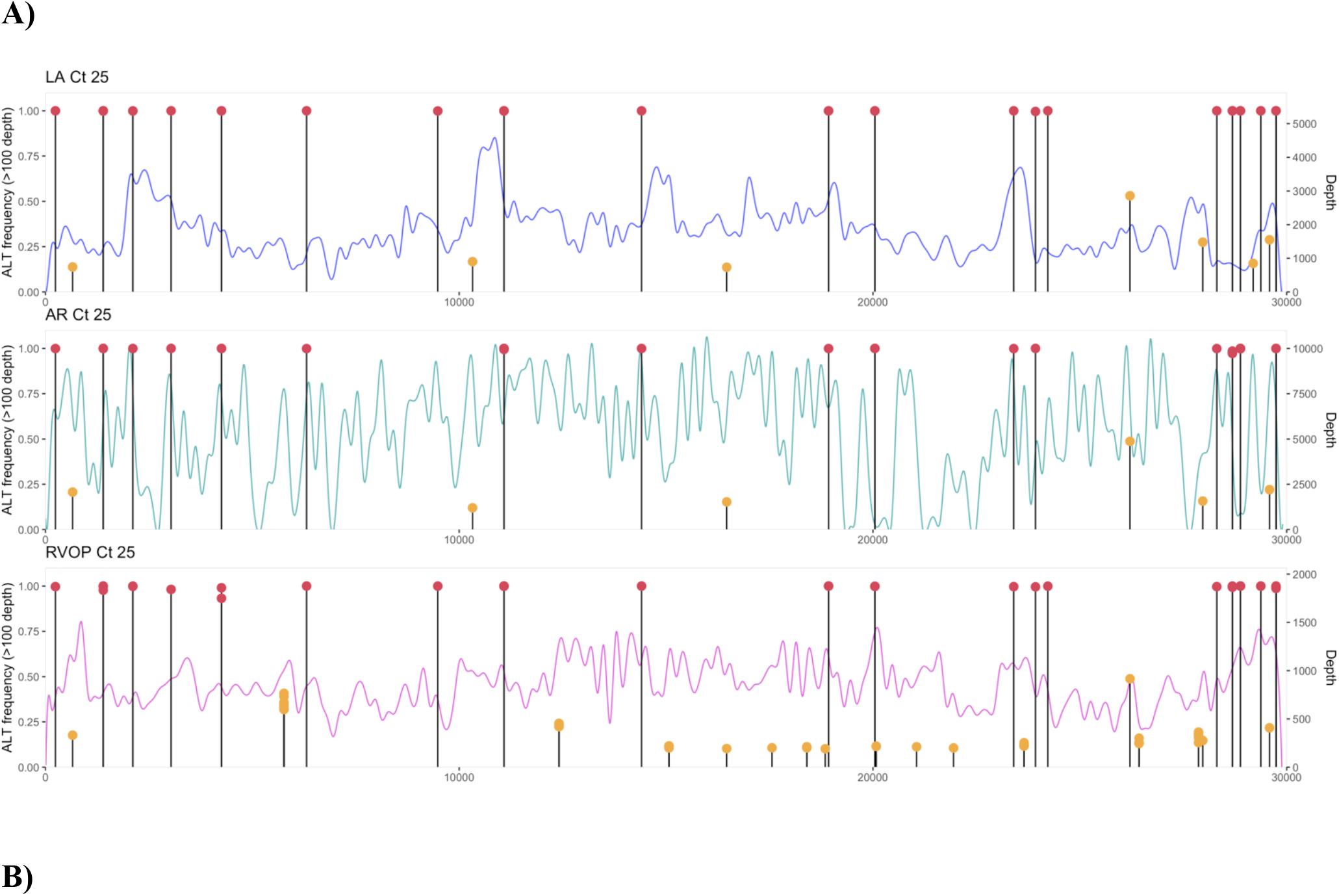
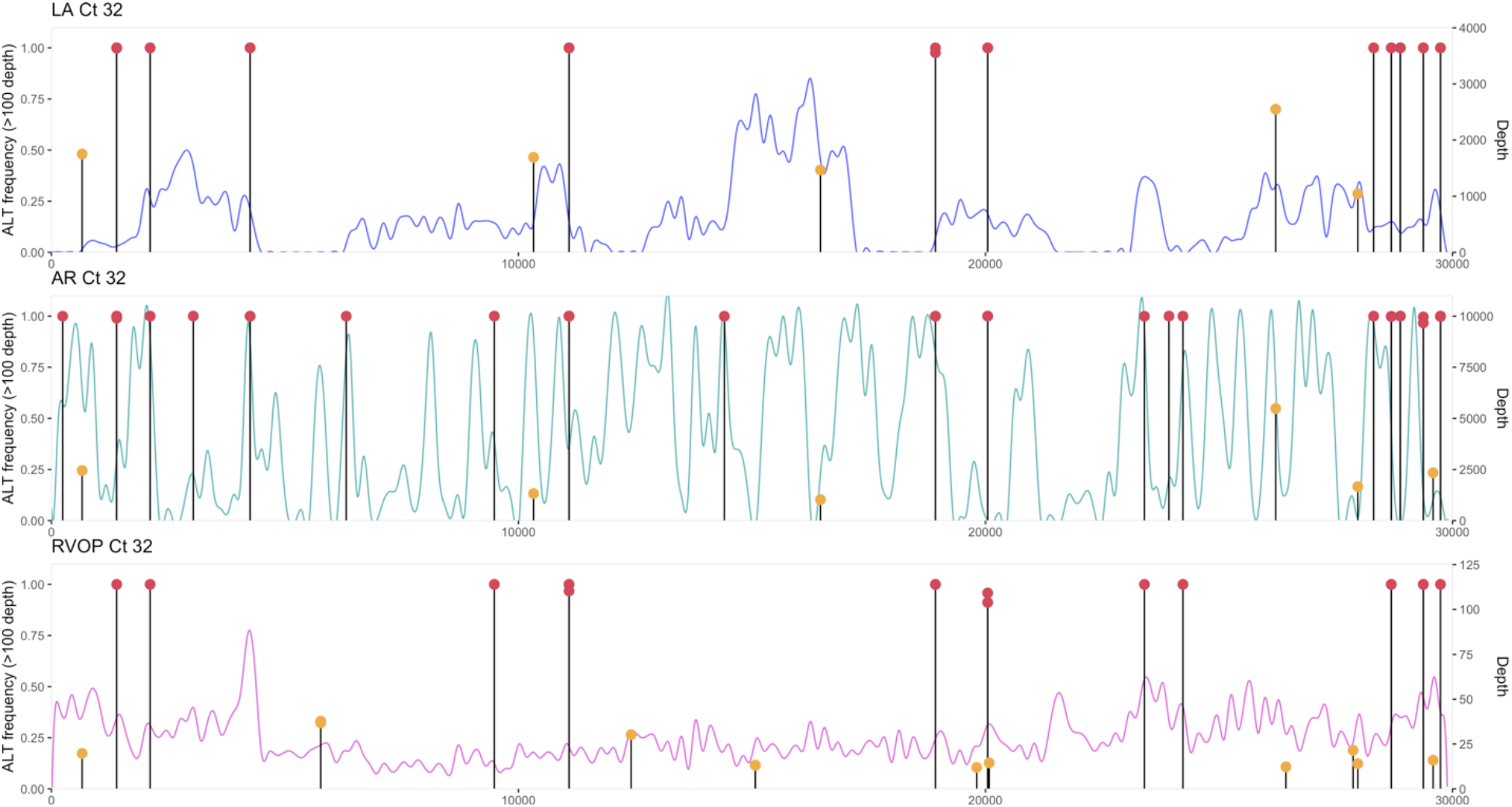
Comparison of read depth for Long-amplification method (LA-blue), ARTIC v3 (AR-green) and the respiratory viral oligo panel (RVOP- pink) at Ct 25 (panel A) and Ct 32 (panel B) and averaged across samples. The depth of coverage is shown on the right axis, while the proportion of reads for single nucleotide polymorphisms (SNPs) and low frequency variant calling is on the left axis. The position of SNPs (red circles) and low frequency variants (orange circles) across the genome are overlaid on top of the both coverage graphs.

